# Huntington’s disease-specific mis-splicing captured by human-mouse intersect-RNA-seq unveils pathogenic effectors and reduced splicing factors

**DOI:** 10.1101/2020.05.11.086017

**Authors:** Ainara Elorza, Yamile Márquez, Jorge R. Cabrera, José Luis Sánchez-Trincado, María Santos-Galindo, Ivó H. Hernández, Juan Ignacio Díaz-Hernández, Ramón García-Escudero, Manuel Irimia, José J. Lucas

## Abstract

Deregulated alternative splicing has been implicated in a wide range of pathologies. Deep RNA-sequencing has revealed global mis-splicing signatures in multiple human diseases; however, for neurodegenerative diseases, these analyses are intrinsically hampered by neuronal loss and neuroinflammation in post-mortem brains. To infer splicing alterations relevant to Huntington’s disease (HD) pathogenesis, here we performed intersect-RNA-seq analyses of human post-mortem striatal tissue and of an early symptomatic mouse model in which neuronal loss and gliosis are not yet present. Together with a human/mouse parallel motif scan analysis, this approach allowed us to identify the shared mis-splicing signature triggered by the HD-causing mutation in both species and to infer upstream deregulated splicing factors. Moreover, we identified a plethora of downstream neurodegeneration-linked effector genes, whose aberrant splicing is associated with decreased protein levels in HD patients and mice. In summary, our intersect-RNA-seq approach unveiled the pathogenic contribution of mis-splicing to HD and could be readily applied to other neurodegenerative diseases for which bona fide animal models are available.

## Introduction

Alternative splicing (AS) of pre-mRNA is the differential processing of introns and exons to generate multiple transcript isoforms from individual genes, thereby increasing molecular diversity. However, when it is not properly executed, AS leads to mis-splicing, which may result in proteins with altered function and stability^1^. A number of splicing factors and other RNA-binding proteins (RBPs) are responsible for proper regulation of AS^2^, and growing evidence has implicated mis-splicing in a range of pathologies such as cancer^3^, muscular dystrophies^4^, autism^5,6^ and neurodegenerative diseases^7^ such as Alzheimer’s disease^8,9^, amyotrophic lateral sclerosis^10,11^ and Huntington’s disease^12-15^. Next-generation RNA sequencing (RNA-seq) has boosted investigation of global mis-splicing in diseased tissue. Nevertheless, while analysis of tumour or dystrophic muscle biopsies can be informative on early stage pathogenic mis-splicing, RNA-seq studies of neurodegenerative diseases using post-mortem brains are confounded by the dramatically altered cellular composition in end-state disease tissue due to neuronal loss and increased gliosis, the latter besides leading to a chronic inflammatory status. These caveats make it difficult to identify the causative molecular alterations that may serve as bases for therapeutic approaches.

Huntington’s disease (HD) is a devastating neurological disorder characterized by prominent motor symptoms and marked atrophy of the nucleus striatum^16^. HD is caused by a polyglutamine (polyQ)-encoding CAG repeat expansion in the Huntingtin (*HTT*) gene^17^. Similar pathogenic CAG mutations in different genes cause multiple dominant spinocerebellar ataxias (SCAs), including SCA-1, -2, -3, -6, -7 and -17^18^ and there is evidence of toxicity being mediated by both the expanded CAG-containing mRNAs and the polyQ-containing proteins^19,20^. The proteins that interact with expanded CAG mRNA include splicing factors such as MBNL1^21^, U2AF2^22^ and SRSF6^15,23^, the latter being also sequestered into the characteristic polyQ inclusion bodies found in HD brains^13^. All this has led to the proposal that splicing alterations may, at least in part, underlie HD^12-15^. In fact, hypothesis-driven studies have identified two mis-splicing events in the etiology of HD. Namely, a retained intron in the *HTT* gene giving rise to a highly toxic exon1-encoded N-terminal truncated form of mutant *Htt*^12^, and elevated inclusion of tau (*MAPT*) exon 10, leading to a pathogenic increase in tau isoforms with four tubulin binding repeats (4R-tau)^13^. However, analysis of global mis-splicing in striatum of HD patients is missing due to the limitations imposed by the altered cellular composition in post-mortem samples discussed above.

To investigate mis-splicing patterns potentially relevant to the onset of HD, here we performed at par RNA-seq analyses of striatum of HD patients and of a highly validated HD transgenic mouse model at an early disease stage, in which motor symptoms are due to circuit dysfunction prior to appearance of neuronal loss and gliosis. Intersection of both analyses yielded a shared mis-splicing signature triggered by the HD causing mutation in striatum of both species (Fig.1). Characterization of this signature allowed us to infer a network of upstream splicing factors deregulated in both species and to identify a plethora of downstream neurodegeneration-linked effector genes whose aberrant splicing led us to corroborate their decreased protein levels in HD striatum.

**Figure 1.**
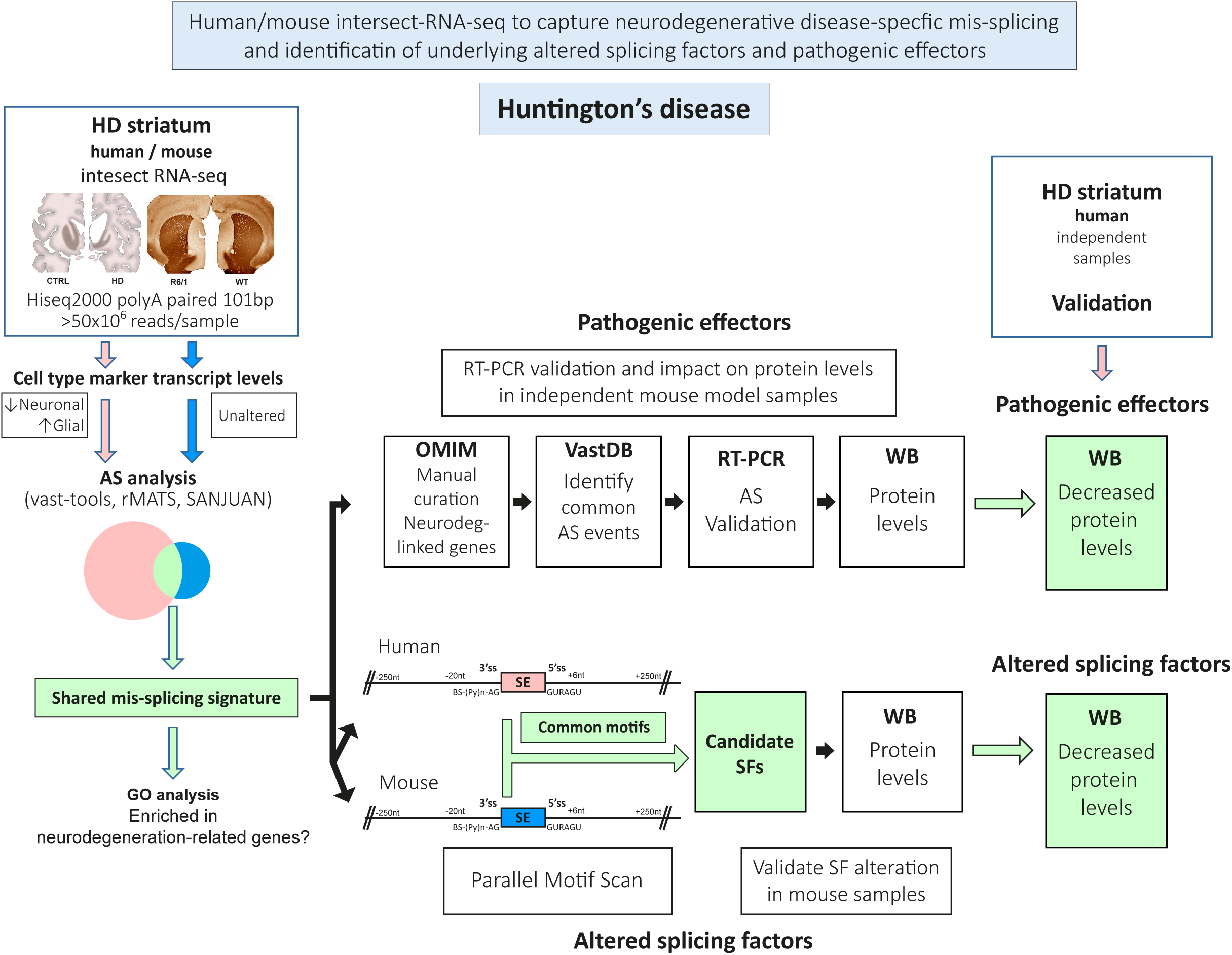
Workflow of the technical approach. Steps for human/mouse intersect-RNA-seq analysis of neurodegenerative disease-specific mis-splicing and identification of underlying altered splicing factors and pathogenic effectors.

## Results

### Intersect-RNA-seq analysis of striatum of HD patients and early symptomatic mice identifies a shared mis-splicing signature affecting movement disorder-related genes

Given the fully penetrant and dominant nature of the HD causing mutation, transgenic animal models that truly recapitulate symptoms and neuropathology have been generated. In particular, mice of the R6/1 model express a transgene driven by the endogenous *Htt* promoter that encodes the highly toxic exon1-encoded N-terminal form of *Htt* with a CAG expansion resulting in a robust, yet slowly-progressing, motor phenotype^24^. At the age of 3.5 months, R6/1 mice in our colony already showed a clear motor coordination deficit (p=5.0×10^−3^; Student’s t-test; Supplementary Fig. 1a), but no significant striatal atrophy or neuropathology, as evidenced by volumetric analysis of the striatum together with stereological neuronal count and immunostaining with markers of gliosis (Supplementary Fig. 1b,c). Given the lack of gliosis and neuronal loss at this stage, we reasoned that a transcriptomic comparison of striatum from this mouse model and from human HD patients could reveal shared HD-specific pathogenic mis-splicing signatures triggered by the HD-mutation in both species, devoid of secondary alteration-related artifacts. Therefore, we performed at par RNA-seq analyses of post-mortem striatum of HD patients (Vonsattel’s grade 3-4) and matching control subjects (n=3), and of striatum of 3.5 month-old R6/1 mice, together with matching controls (n=3). Between 55 and 102 million reads (101-bp, pair-ended) were generated per sample (Fig. 2a). Differential gene expression analysis confirmed no alterations in the levels of neuronal and glial marker genes in the R6/1 RNA-seq samples (Fig. 2b). In contrast, as expected from the marked neuronal loss and gliosis of human HD post-mortem striatum^25^, we observed a strong decrease in the relative expression of neuronal marker genes (p=8.0×10^−5^; Student’s t-test) and an increase in glial-marker genes, particularly those of astrocytes (p=0.018; Student’s t-test) and microglia (p=4.2×10^−3^; Student’s t-test) (Fig. 2b).

**Figure 2.**
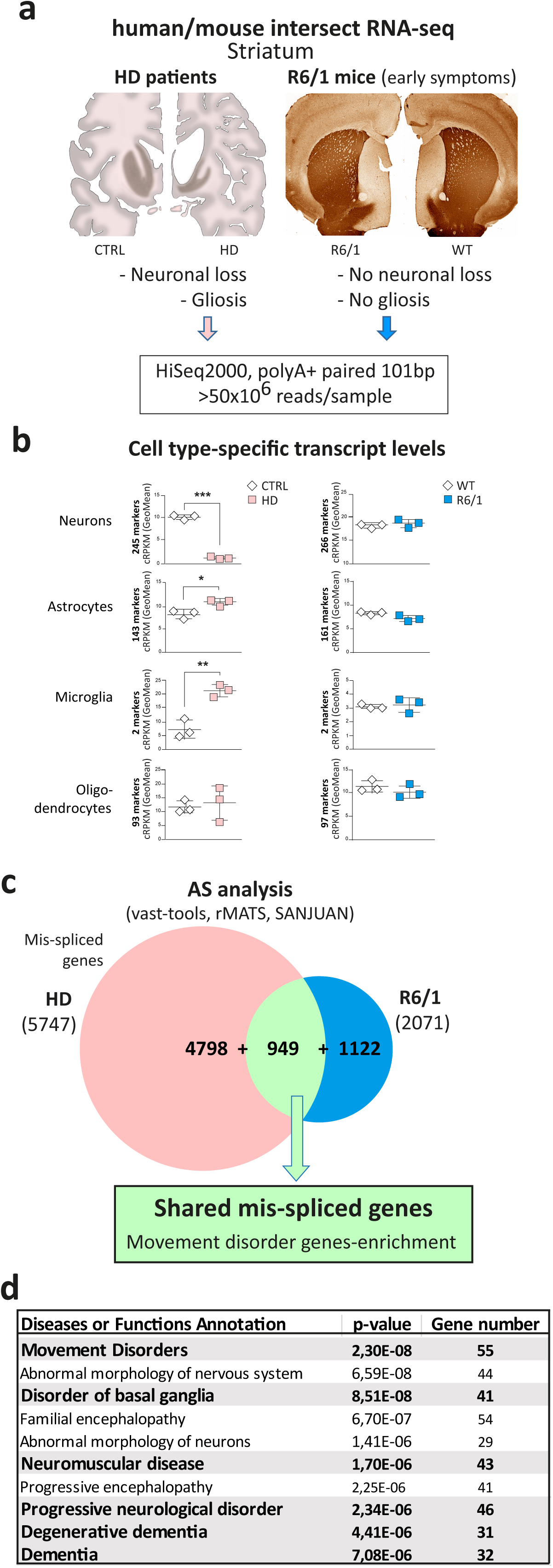
Intersect-RNA-seq analysis of striatum of HD patients and early symptomatic R6/1 mice. **(a)** For at par RNA-seq analysis, polyA+ RNA was prepared from post-mortem striatum of HD patients (Vonsattel’s grade 3-4) and matching control subjects (*n*=3), and from striatum of 3.5 month-old R6/1 mice, together with matching controls (*n*=3). The schematic drawings on the left depict coronal sections of human control and HD patient brain showing striatal atrophy and micrographs on the right show representative images of DARPP32 immunostaining of non-atrophied striatum in coronal sections of 3.5 month-old R6/1 mice as compared to WT mice. **(b)** Analysis of levels of cell type-specific transcripts in human (left, *n*=3) and mouse (right, *n*=3) striatal RNA-seq samples using geometric mean of the cRPKM values of cell type marker genes of neurons, astrocytes, microglia and oligodendrocytes (Student’s t-test; \**P* < 0.05, *\*\* P < 0*.*01*, \*\*\**P* < 0.001). **(c)** Splicing was analyzed with *vast-tools*, rMATs and SANJUAN software and the Venn diagram shows the number of mis-spliced genes in HD patients (5,747) and R6/1 mice (2,071) with respect to controls, as well as the intersect of 949 mis-spliced genes common to both species (p = 6.1×10-3, two-sided Fisher’s Exact test using only one-to-one orthologs with sufficient read coverage in both species as background; N = 12882). **(d)** Ingenuity Pathways Analysis (IPA) of the 342 human/mice shared mis-spliced genes detected by single-program RNA-seq analysis shown in Supplementary Table 10.

Next, we compared AS between HD and control samples in both species using three complementary software packages (*vast-tools*, rMATS and SANJUAN; see Methods for details). Combining the results obtained from each tool produced a list of 5,747 genes with at least one differentially spliced AS event in HD patients respect to control subjects (Fig. 2c, Supplementary Fig. 2 and Supplementary Table 1). Since isoform usage differs widely across cell types^26,27^ and is regulated upon inflammation^28^, a substantial fraction of the observed AS changes are expected to be due to the altered neuronal/glia cellular composition and/or to the pro-inflammatory signals in post-mortem HD tissue. In line with this, the number of differentially spliced genes in R6/1 mice respect to controls was much smaller (2,071 genes; Fig. 2c, Supplementary Fig. 2 and Supplementary Table 2). Remarkably, a total of 949 one-to-one orthologs were differentially spliced in both human and mouse (p=6.12×10^−3^, two-sided Fisher’s Exact test; Fig. 2c and Supplementary Tables 3-10). This set of genes are expected to reflect AS alterations that are mainly due to the toxicity of the HD mutation independently of altered cellular content-associated artifacts in the human samples. Accordingly, this shared mis-splicing signature represents an important fraction of all mis-spliced genes in the mouse model (45.8%) but only a minority of those observed in human tissue (16.5%). Interestingly, Gene Ontology analysis revealed that common mis-spliced genes were strongly enriched for functions related to basal ganglia and movement disorders (including HD)(Fig. 2d and Supplementary Table 11), suggesting that these shared AS alterations in striatum are a plausible contributor to HD pathogenesis.

### Mis-splicing of neurodegeneration-linked genes correlates with reduced protein levels

To further explore the pathogenic relevance of mis-spliced genes, we next focused on a subset of manually curated genes with differential AS in both species and whose mutations cause monogenic forms of neurodegeneration in humans: *CCDC88C* (Coiled-coil domain containing 88C linked to SCA40 -spinocerebellar ataxia 40-, OMIM #616053), *KCTD17* (Potassium channel tetramerization domain containing 17 linked to DYT26 -myoclonic dystonia-26-, OMIM #616398), *SYNJ1* (Synaptojanin 1 linked to PARK20 -early onset Parkinson disease-20-, OMIM #615530), *VPS13C* (Vacuolar protein sorting 13 homolog C linked to PARK23 -early onset Parkinson disease-23-, OMIM #616840), *TRPM7* (Transient receptor potential cation channel subfamily M member 7 linked to ALSP/DC -amyotrophic lateral sclerosis-parkinsonism/dementia complex-, OMIM #105500) and *SLC9A5* (Solute carrier family 9 member A5 also known as Na(+)/H(+) exchanger 5 that has been suggested to be linked to EKD2 -episodic kinesigenic dyskinesia 2-, as it maps to the 16q13-q22.1 candidate region^29^) (Fig. 3a). Using VastDB^30^, we identified the orthologous AS events that were differentially spliced in both species, and which corresponded to skipped exons (SE) in all cases except for *TRPM7*, which had a differentially retained intron (RI) (Fig. 3b). All these mis-spliced events detected by RNA-seq were validated by RT-PCR assays using striatal mRNA from an independent set of R6/1 and wild-type mice (n=5; Fig. 3c).

**Figure 3.**
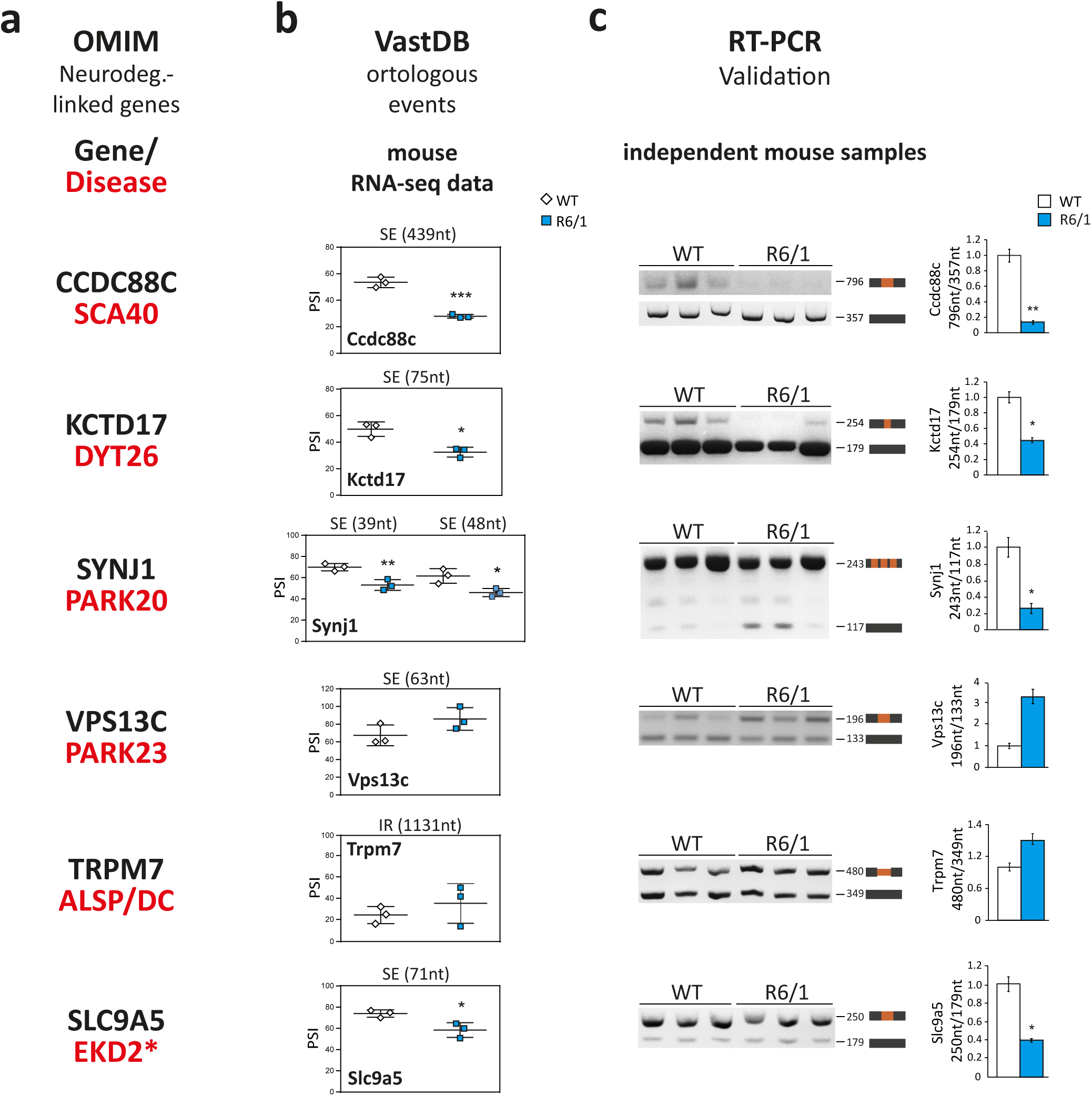
Mis-splicing of neurodegeneration-causing genes. **(a)** The OMIM database was used to select, among the genes that are mis-spliced in both human and mouse striatum, genes whose mutations in humans cause monogenic forms of neurodegeneration. **(b)** Using VastDB^30^, the orthologous alternative spliced events differentially processed in both species were identified and the PSI values of the mis-spliced events in each gene are shown in 3 WT vs 3 R6/1 samples (according to the RNA-seq *vast-tools* analysis). **(c)** RT-PCR assay and quantification of mis-spliced events in striatal RNA from an independent set of WT and R6/1 mice (*n*=5). (Student’s t-test; \**P* < 0.05, *\*\* P < 0*.*01*). Data represent mean ± SEM.

Next, we tested whether the splicing alterations observed in striatum of R6/1 mice in these neurodegeneration-associated genes correlated with changes in their protein levels. Western blot analyses revealed a significant decrease in protein levels for all of them except for Vps13c, which also showed a trend of reduced protein levels respect to the control samples (Fig. 4). Since both recessive mutations and decreased protein levels may result in decreased protein function, these data suggest that the observed global mis-splicing in striatum of HD patients and mice may have a pathogenic effect by diminishing protein levels of multiple neurodegeneration-linked genes.

**Figure 4.**
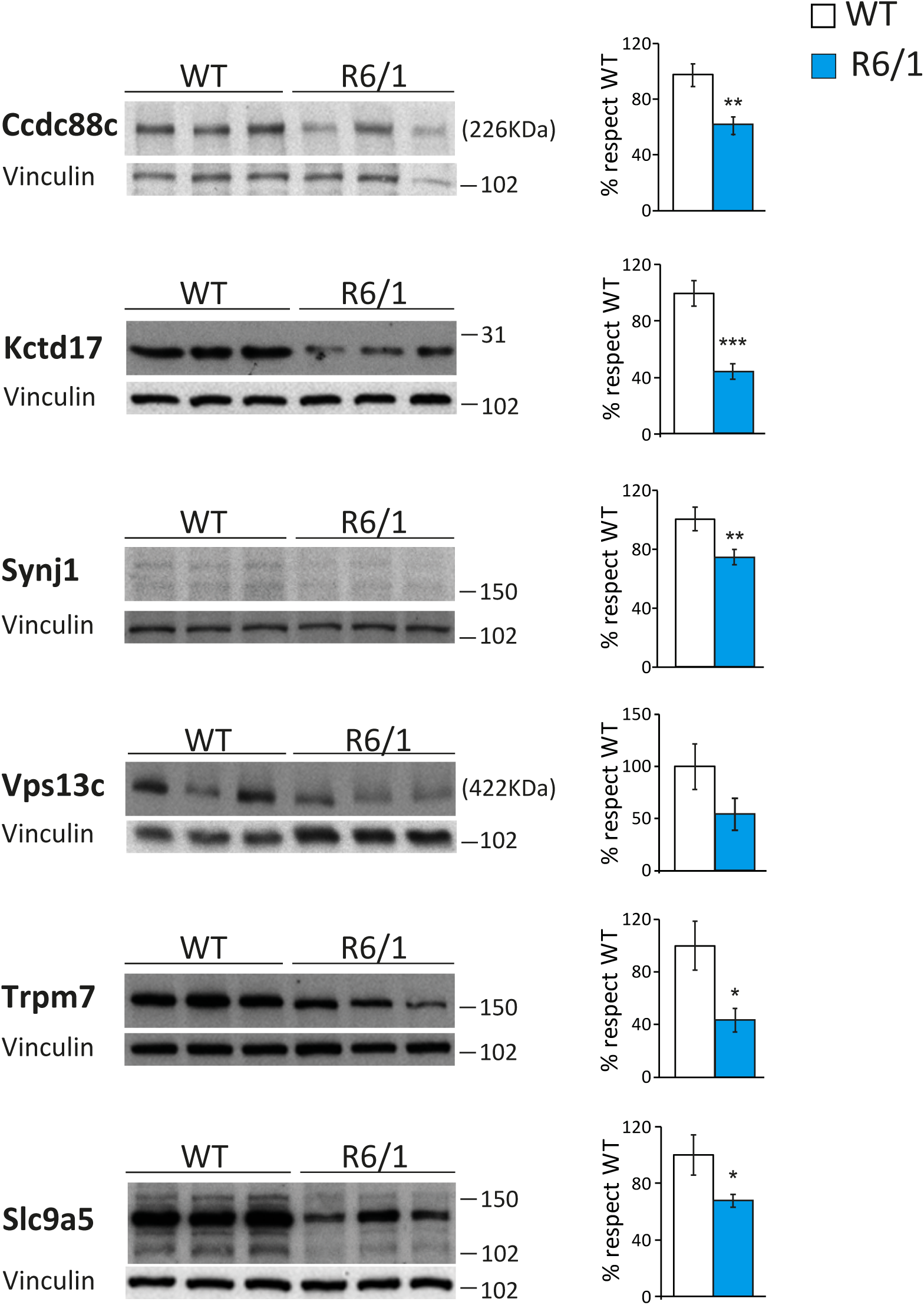
Decreased protein levels of neurodegeneration-causing genes in striatum of R6/1 mice. Protein levels in striatum of WT and R6/1 mice and quantifications normalized with Vinculin (*n*=7-12). (Student’s t-test; \**P* < 0.05, *\*\* P < 0*.*01*, \*\*\**P* < 0.001). Data represent mean ± SEM.

### Parallel RBP motif enrichment analysis reveals splicing factors altered in HD striatum

We next looked for potential RBPs that may be responsible for the shared mis-splicing HD signature. For this purpose, we searched for known binding motifs for RBPs^31^ inside and around orthologous exons that were mis-regulated in both species. We reasoned that motifs associated with relevant RBPs would be similarly enriched in both exons sets. Since *vast-tools* directly provides exon-based homology information between the two species, we focused on 73 orthologous alternative exons detected by this software as mis-spliced in both human and mouse (|ΔPSI| ≥ 15) (Supplementary Table 12). As a control gene set, we used 914 alternatively spliced exons whose usage was not altered in HD and R6/1 samples respect to controls (|ΔPSI| < 5) (Fig. 5a). Motifs associated with six RBP families showed significant enrichments in the equivalent positions in both species: binding motifs of TIA1, U2AF2, HNRNPC and PTBP were enriched in the upstream intronic sequences and of RBFOX and ELAVL in downstream intronic sequences (Fig. 5b).

**Figure 5.**
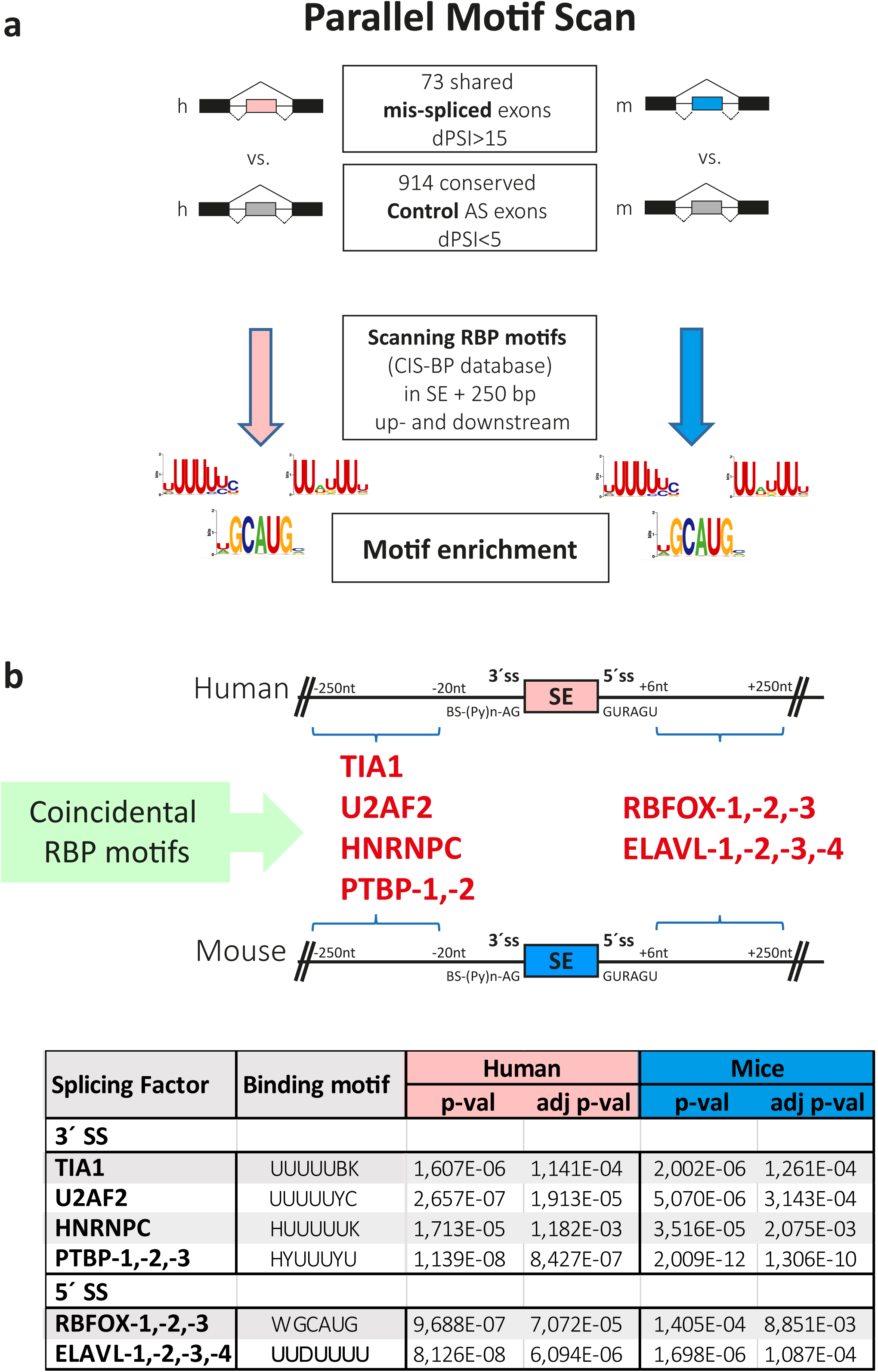
Human-mouse parallel RBP motif scan analysis in mis-regulated alternatively spliced events identifies enrichment of several splicing factor binding motifs. **(a)** Schematic representation of scan RBP motif analysis performed in the 73 differentially included alternative exons detected with *vast-tools* and conserved in human (h) and mice (m). **(b)** Significantly enriched binding sites of splicing factors in 250 intronic base pairs adjacent to differentially skipped exons in HD and R6/1 samples.

Western blot analysis of these RBPs in striatum of R6/1 and control mice revealed a significant decrease in protein levels for nine different members belonging to five of the six identified RBP families. Specifically, we found decreased levels for TIA1 (71%, p=9.2×10^−4^; Student’s t-test), U2AF2 (26%, p=2.2×10^−3^; Student’s t-test), RBFOX1 (28% p=0.016; Student’s t-test), RBFOX2 (46%, p=8.5×10^−3^; Student’s t-test), RBFOX3 (60%, p=1.3×10^−5^; Student’s t-test), ELAVL4 (76%, p=1.5×10^−4^; Student’s t-test), ELAVL2 (55%, p=2.3×10^−5^; Student’s t-test), ELAVL1 (30%, p=2.9×10^−4^; Student’s t-test) and HNRNPC (66%, p=1.9×10^−3^; Student’s t-test) (Fig. 6). In contrast, the protein levels of ELAVL3 did not show any change in R6/1 striatum, neither did the two paralogs of the PTBP family, which were preferentially expressed in astrocytes^32^ (Supplementary Fig. 3).

**Figure 6.**
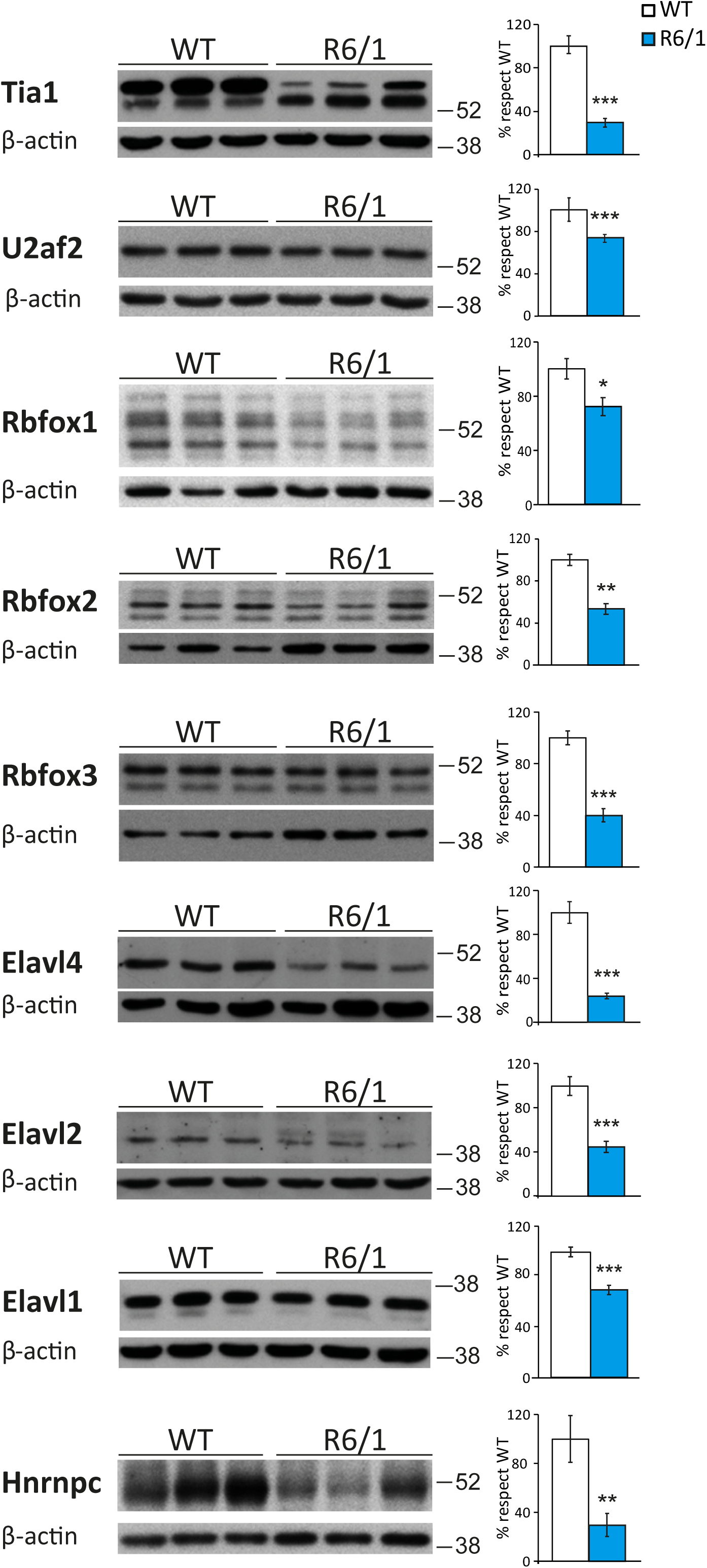
Decreased protein levels of splicing factors in striatum of HD mouse model. Protein levels in striatum of WT and R6/1 mice and quantifications normalized with β-actin (*n*=7-12). (Student’s t-test; \**P* < 0.05, *\*\* P < 0*.*01*, \*\*\**P* < 0.001). Data represent mean ± SEM.

### Validation of altered splicing factors and pathogenic downstream effectors in striatum of HD patients

Finally, we assessed whether the protein-level alterations observed in R6/1 striatum for both splicing factors and neurodegenerative-associated genes reflected matching alterations in the striatum of HD patients, as expected from our intersect-RNA-seq analysis. Regarding the splicing factors, we found decreased protein levels for TIA1 (91%, p=0.036; Student’s t-test), U2AF2 (65%, p=0.045; Student’s t-test), RBFOX1 (76%, p=5.3×10^−4^; Student’s t-test), RBFOX2 (46%, p=4.6×10^−3^; Student’s t-test), RBFOX3 (96%, p=3.3×10^−3^; Student’s t-test) and ELAVL2 (62%, p=1.8×10^−4^; Student’s t-test). Besides, a similar trend for decreased protein levels (53%, p=0.17; Student’s t-test) was also observed for HNRNPC (Fig. 7a). For the subset of neurodegeneration-linked mis-spliced genes, we observed a sharp decrease in protein levels for virtually all of them (Fig. 7b). More precisely, for CCDC88C we found a 90% decrease in a low molecular weight form (p=3.2×10^−5^; Student’s t-test) and a trend for decrease in the canonical 228 kDa form (60%, p=0.081; Student’s t-test), for KCTD17 we observed a 58% decrease (p=1.7×10^−3^; Student’s t-test) in the monomeric (36 kDa) form and a 59% decrease (p=0.048; Student’s t-test) in the pentameric form (data not shown), and we also observed decreased levels of SYNJ1 (56%, p=0.021; Student’s t-test), TRPM7 (82%, p=6.0×10^−3^; Student’s t-test), VPS13C (69%, p=0.012; Student’s t-test) and SLC9A5 (83%, p=1.0×10^−3^; Student’s t-test) (Fig. 7b). Altogether, these results indicate that the mis-splicing HD signature and the candidate splicing factors identified by our intersect-RNA-seq and parallel motif scan analyses show high validation rates also at the protein level in striatum of HD patients.

**Figure 7.**
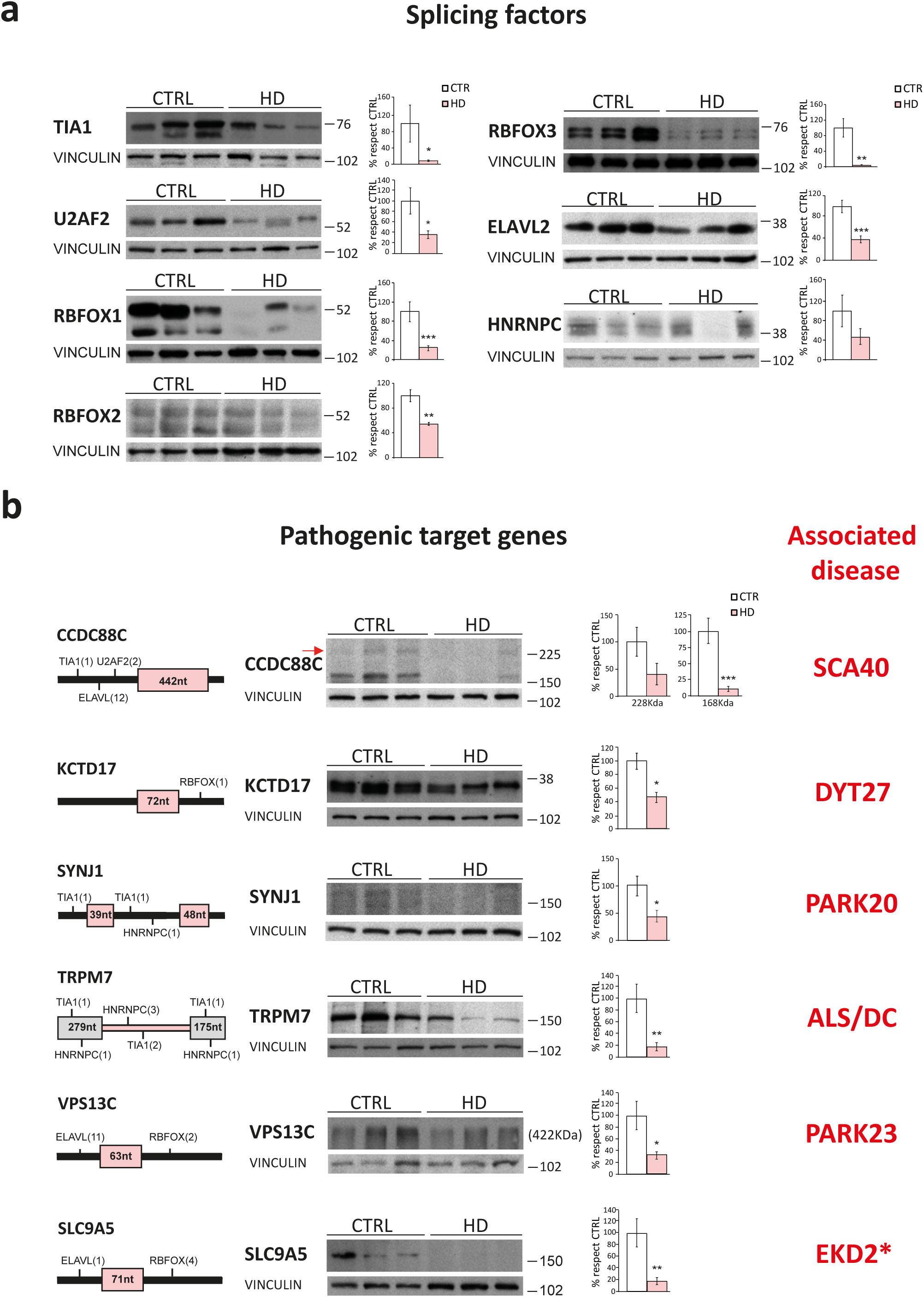
Alteration of splicing factors and neurodegeneration-causing genes protein levels in striatum of HD patients. **(a)** Representative western blots of protein levels of splicing factors in striatum of HD patients and controls and quantification normalized with Vinculin (*n*=7-13). (Student’s t-test; \**P* < 0.05, *\*\* P < 0*.*01*, \*\*\**P* < 0.001). Data represent mean ± SEM. **(b)** Localization of binding motifs of splicing factors enriched in the proximity of the mis-spliced events in each pathogenic gene and their protein levels in control and HD striatum. Histograms show quantification of protein levels normalized with Vinculin (*n*=7-13). (Student’s t-test; \**P* < 0.05, *\*\* P < 0*.*01*, \*\*\**P* < 0.001). Data represent mean ± SEM.

## Discussion

Here we report a novel approach to identify potentially pathological signatures of mis-splicing in HD patients overcoming the artifacts associated with altered cell type composition and neuroinflammation in human post-mortem brain tissues. We performed at par RNA-seq analysis of striatal tissue from a mouse model of HD at an early symptomatic stage and from human post-mortem HD brain, whose intersection revealed a shared HD-specific mis-splicing signature affecting 949 one-to-one orthologous genes. These included a subset of genes that had been previously involved in neurodegenerative movement disorders and that we showed to have reduced protein levels in the mouse model as well as in HD patients. In addition, by human-mouse parallel complementary motif searches on common mis-spliced events, we inferred a network of candidate upstream splicing factors with reduced protein levels in both species.

A previous report also addressed global mis-splicing in human post-mortem HD tissue by RNA-seq analysis^14^. However, instead of studying the striatum, which is the brain region primarily and most affected in the disease, this study focused on BA4 motor cortex, which shows a less marked atrophy in post-mortem tissue. Nevertheless, it is well documented that the cortex also degenerates early in the course of disease^33^, and therefore is likely affected by similar caveats associated to neurodegeneration and neuroinflammation. In line with this, Lin and co-workers detected enrichment of 15 splicing factor motifs in their set of differential AS events, among which only PTBP1 overlaps with the splicing factors detected in our analysis. However, PTBP1 is preferentially expressed in astrocytes^32^ and is not altered at the protein level neither in human or mouse HD striatum, unlike the other five families of splicing factors that we detected through our parallel motif scan. Besides, out of the nine candidate splicing factors that show altered mRNA levels in Lin’s work, the majority also change in human striatum according to our RNA-seq data, but only one is altered in striatum of early symptomatic mice. Overall, these observations point to neurodegeneration-associated confounding factors and strengthen the notion that, by refining mis-splicing analysis by intersecting results from human patients and an early mouse model, neurodegeneration- and neuroinflammation-associated artifacts are cleared out in our study.

Eight additional neurological disorders including the spinal and bulbar muscular atrophy (SBMA) and several SCAs are caused by expanded CAG repeats in coding sequences of different genes that give rise to expanded polyQ containing proteins^18^. Given the similarity in the causing mutations, the molecular mechanisms in HD pathogenesis -including splicing alterations-, are likely contributors to the other CAG repeat polyglutaminopathies. Most of these diseases show maximal affectation in cerebellum and the analysis of global mis-splicing in post-mortem cerebellar patient tissue for these diseases poses the same challenges as those addressed here for HD striatum. Since there are also good animal models for these triplet repeat disorders^34^, at par early mouse model RNA-seq to identify the intersecting mis-splicing signature and parallel motif scans to infer associated splicing regulators will be useful to determine whether similar AS-related pathogenic mechanisms underlie the different CAG trinucleotide repeat disorders. Similarly, the human-mouse intersect-RNA-seq approach might be applied to any neurodegenerative disease for which animal models with construct validity are available and for which alterations in RNA processing are suspected as a pathogenic mechanism. These include the amyotrophic lateral sclerosis/frontotemporal dementia disease continuum (ALS/FTD)^35^, particularly the forms associated to mutations in genes encoding RBPs (*TDP43* and *FUS*) or affecting formation of RNA foci (*C9orf72*)^36^ for which bona fide animal models exist^1^.

Pathological expanded CAG-containing mRNAs adopt a hairpin conformation^37^ and some of the altered splicing factors detected here are likely to be affected by direct and aberrant interaction with expanded CAG mRNA. This is in fact the case of U2AF2^15,22^ and HNRNPC^15^. Both U2AF2 and HNRNPC bind to U-rich motifs, as also do TIA1 and the ELAVL family, all identified here as mis-regulated in human HD patients and the mouse model. Since they compete for similar target mRNAs and may be regulated by common interactors, it is conceivable that deregulation of TIA1 and ELAVL might be secondary to U2AF2 and HNRNPC alterations driven by their interaction with expanded CAG repeats. The RBFOX family is the only one altered in our study that does not bind U-rich motifs, as it binds the GCAUG consensus^38^. However, RBFOX splicing factors also act through a multiprotein complex called LASR (Large Assembly of Splicing Regulators) that includes HNRNPC^39^ and it is conceivable that sequestration of HNRNPC by expanded CAG-HTT exon1 RNA^15^ might subsequently affect RBFOX binding and stability.

In summary, by intersect-RNA-seq analysis of at par-processed early mouse model and human post-mortem brain tissue, we have identified a HD-specific mis-splicing signature that allowed us to infer altered upstream splicing factors and downstream pathogenic effectors that can become new therapeutic targets for HD. Moreover, this approach can be applied to investigate the potential relevance of altered AS in other neurodegenerative diseases.

## Acknowledgements

This work was supported by CIBERNED-ISCIII collaborative grants PI2015-2/06-3 and PI2018/06-1 ; by grants : SAF2015-65371-R (MINECO/AEI/FEDER, UE) ; RTI2018-096322-B-I00 (MCIU/AEI/FEDER, UE) from Spanish Ministry of Economy and Competitiveness/Ministry of Science, Innovation and Universities (MINECO/MICINN) to JJL and BFU2017-89201-P (MINECO/AEI/FEDER, UE) to MI; PI18/00263 from the Instituto de Salud Carlos III (Ministry of Economy, Industry and Competitiveness) co-funded by the European Regional Development Fund-to RG-E; institutional grant from Fundación Ramón Areces to CBMSO ; Fundación BBVA to JJL; and by the European Research Council under the European Union’s Horizon 2020 research and innovation program ERC-StG-LS2-637591 to MI. Human tissue was obtained from Institute of Neuropathology (HUB-ICO-IDIBELL) Brain Bank, the Neurological Tissue Bank of the IDIBAPS Biobank, the Banco de Tejidos Fundación CIEN, and the Netherlands Brain Bank. We thank Juan Valcárcel for scientific advice. We also thank excellent technical assistance by Miriam Lucas and by the following core facilities: CBMSO-Genomics & Massive Sequencing, CBMSO-Animal Facility and the CETA-CIEMAT computing center. We also thank Alvaro Lucas for help with illustrations.

## Materials and Methods

### Human brain tissue samples

Brain specimens used in this study from striatum of HD patients and controls (CTRL) were provided by Institute of Neuropathology Brain Bank (HUB-ICO-IDIBELL, Hospitalet de Llobregat, Spain), the Neurological Tissue Bank of the IDIBAPS Biobank (Barcelona, Spain), the Banco de Tejidos Fundación Cien (BT-CIEN, Madrid, Spain) and the Netherlands Brain Bank (Amsterdam, The Netherlands). Written informed consent for brain removal after death for diagnostic and research purposes was obtained from brain donors and/or next of kin. Procedures, information and consent forms have been approved by the Bioethics Subcommittee of Consejo Superior de Investigaciones Científicas (CSIC, Madrid, Spain). The post-mortem interval in tissue processing was between 5:00 and 7:45 h for RNA-seq analysis and between 05:00 and 23:30 h for Western blot analyses. The neuropathological examination in HD cases revealed a diagnosis of HD grade 3-4 following Vonsattel’s criteria^25^.

### Mice

R6/1 transgenic mice for the human exon-1-Htt gene^24^ in B6CBAF1 background were bred and housed at the Centro de Biología Molecular Severo Ochoa animal facility. Mice were housed four per cage with food and water available *ad libitum* and maintained in a temperature-controlled environment on a 12/12 h light-dark cycle with light onset at 08:00. Animal housing and maintenance protocols followed the guidelines of Council of Europe Convention ETS123. Animal experiments were performed under protocols (PROEX293/15) approved by the Centro de Biología Molecular Severo Ochoa Institutional Animal Care and Utilization Committee (Comité de Ética de Experimentación Animal del CBM, CEEA-CBM), Madrid, Spain.

### Rotarod test

Motor coordination was assessed with an accelerating rotarod apparatus (Ugo Basile, Comerio, Italy). Mice were trained during two consecutive days, the first day: 4 trials at fixed 4 rpm for 1 min each, the second day: 4 trials for 2 min (the first minute at 4 rpm and the second minute at 8 rpm). On the third day rotarod was set to accelerate from 4 to 40 rpm over 5 min and mice were tested in 4 trials. The latency to fall from the rotarod was measured as a mean of the 4 accelerating trials.

### Tissue preparation for staining

WT and R6/1 mice euthanasia was performed using CO2. Brain was quickly extracted and the left hemisphere was immersed in 4% paraformaldehyde overnight. After profuse washing in PBS, hemispheres were immersed in sucrose 30% in PBS for at least 72h and then included in OCT (Optimum Cutting Temperature compound, Tissue-Tek, Sakura Finetek Europe, ref. 4583), frozen and stored at -80°C until use. Mouse sagittal and coronal sections (30 µm thick) were sequentially cut on a cryostat (Thermo Scientific), collected and stored free floating in glycol-containing solution (30% glycerol, 30% ethylene glycol in 0.02M phosphate buffer) at -20°C.

### Immunohistochemistry

Sagittal sections were first washed in PBS and then immersed in 0.3% H2O2 in PBS for 45 min to quench endogenous peroxidase activity. After PBS-washes, sections were immersed for 1 h in blocking solution (PBS containing 0.5% FBS, 0.3% Triton X-100 and 1% BSA) and incubated overnight at 4 °C with anti-DARPP32 (1:5000, Chemicon, AB1656) or anti-IBA1 (1:500, Wako, 019-19741) diluted in blocking solution. After washing, brain sections were incubated first with biotinylated goat anti-rabbit secondary antibody and then with avidin-biotin complex using the Elite Vectastain kit (Vector Laboratories, PK-6101). Chromogen reactions were performed with diaminobenzidine (SIGMAFAST DAB, Sigma, D4293) for 10 min. Mouse sections were mounted on glass slides and coverslipped with Mowiol (Calbiochem, Cat. 475904). Images were captured using an Olympus BX41 microscope with an Olympus camera DP-70 (Olympus Denmark A/S).

### Striatal volumetry

Coronal sections (30 µm thick) were cut on a cryostat and every sixth section was counterstained with toluidine blue pH 4.0 (1 g/l Toluidine Blue, Sigma, 198161 in 0.8 M glacial acetic acid). Digital images were captured at a 2.5x magnification (Canon EOS 450D digital camera) and striatal areas from 19-22 sections for each animal were calculated by means of the ImageJ software^40^. Considering a separation of 180 μm between each section, total structure volume in each mouse was calculated.

### Stereology

Coronal sections (30-μm thick) counterstained with toluidine blue pH 4.0 (1g/l toluidine blue (Sigma, 198161), 0.8 M glacial acetic acid) from the volumetric analysis were used. Sections containing striatum were selected and the 15 most central sections were analyzed. One randomly selected 60 µm × 60 µm optical dissector at 60× magnification with an Olympus BX41 microscope with an Olympus camera DP-70 (Olympus Denmark A/S) was analyzed in each section. Total neuronal cell number per dissector was assessed by a researcher blind to genotype. Striatal neuronal cell density was calculated and compared for WT (*n*=3) and R6/1 mice (*n*=3).

### RNA sequencing

Total RNA was isolated using the Maxwell 16 LEV simplyRNA Tissue Kit (Promega, AS1280) and quantified by Qubit® RNA BR Assay kit (Thermo Fisher Scientific) and the RNA integrity number (RIN) was estimated by using RNA 6000 Nano Bioanalyzer 2100 Assay (Agilent). The RNA-seq libraries were prepared with KAPA Stranded mRNA-Seq Illumina® Platforms Kit (Roche) following the manufacturer’s recommendations. Briefly, 500 ng of total RNA was used as the input material, the poly-A fraction was enriched with oligo-dT magnetic beads and the mRNA was fragmented. The strand specificity was achieved during the second strand synthesis performed in the presence of dUTP instead of dTTP. The blunt-ended double stranded cDNA was 3’adenylated and Illumina platform compatible adaptors with unique dual indexes and unique molecular identifiers (Integrated DNA Technologies) were ligated. The ligation product was enriched with 15 PCR cycles and the final library was validated on an Agilent 2100 Bioanalyzer with the DNA 7500 assay. The libraries were sequenced on HiSeq 2500 (Illumina, Inc) with a read length of 2×101bp using HiSeq 4000 SBS kit in a fraction of a HiSeq 4000 PE Cluster kit sequencing flow cell lane generating 55-102 million paired-end reads per sample. Image analysis, base calling and quality scoring of the run were processed using the manufacturer’s software Real Time Analysis (RTA 2.7.7).

### RNA-seq analysis

Differentially expressed genes between the biological groups were analyzed using *vast-tools* v1.1.0 *compare_expr* with default parameters^30^. To verify correct clustering according to genotype, hierarchical clustering of cRPKM expression values was performed using “hclust” and “heatmap.2” R packages over significantly changed genes in HD vs CTRL and R6/1 vs WT mice (Benjamini-Hochberg correction method). Cell-type-marker genes (neurons, astrocytes, oligodendrocytes and microglia) as described previously^14^ were analyzed by calculating their geometric mean of cRPKM values for each cell type excluding those genes with cRPKM null value in at least one sample and data were plotted and analyzed using GraphPad software (La Jolla, CA).

To identify differential alternative splicing events between the two sample groups per species we used three complementary software: (1) *vast-tools* v1.1.0, using human (hg19; vastdb.hsa.13.11.15) or mouse (mm9; vastdb.mmu.13.11.15) junction libraries for the *align* and *combine* modules^30^. Differential splicing analysis was done using the module *compare* with default parameters (|ΔPSI| ≥ 15 and a minimum ΔPSI of 5 in the same direction among all replicates). (2) rMATS, v3.0.8^41^, utilizing TopHat2^30^ to align fastq reads against the GRCh38.p2 (dec. 2014) and GRCm38.p3 (jan. 2012) genomes together with a custom splice-junction library. For differential splicing analyses, default parameters were used (FDR < 5%). (3) SANJUAN v1.0-beta, which detects de-novo splicing junctions and AS events (https://github.com/ppapasaikas/SANJUAN). For differential splicing analyses, medium confidence level of constraints on differential splicing junctions and |ΔPSI| ≥ 15 were used. The relative contribution of each tool to the detection of mis-spliced genes is shown in Supplementary Fig. 2 and Supplementary Tables 3-8.

The entire RNA-seq dataset will be available at the Gene Expression Omnibus (GEO) database with the accession number XXXXX.

### Ingenuity Pathway Analysis (IPA) analysis

Genes mis-spliced in both human and mouse were annotated to diseases and functions using Ingenuity Pathway Analysis (IPA) software (http://www.ingenuity.com). The analysis was performed in October 2019 and the results were filtered for neurological diseases related functions.

### Semiquantitative real-time PCR validation

Selected events were evaluated with semiquantitative RT–PCR. Total RNA (500 ng) was reverse-transcribed using Invitrogen Superscript IV reverse transcriptase and cDNA (50 ng) was amplified with gene/exon-specific primers. PCR products were resolved on 2% high-resolution Metaphor agarose gels (Lonza).

### Primers

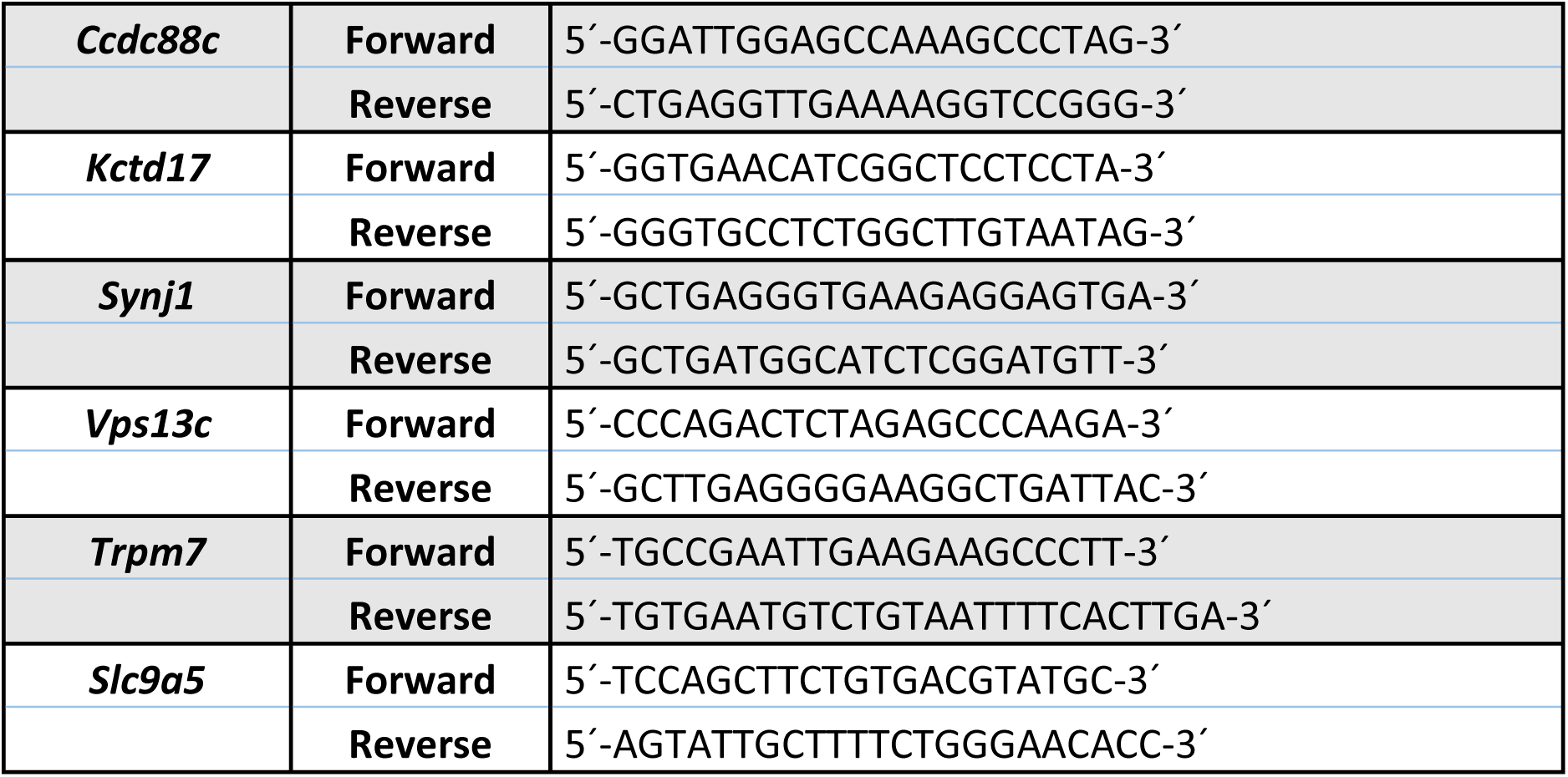

#### Western Blot

Samples from human brain were stored at -80°C and ground with a mortar in a frozen environment with liquid nitrogen to prevent thawing of the samples, resulting in tissue powder. Mouse brains were quickly dissected on an ice-cold plate and the different structures stored at -80°C. Human and mouse protein extracts were prepared by homogenizing brain structures in ice-cold extraction buffer (20 mM HEPES pH 7.4, 100 mM NaCl, 20 mM NaF, 1% Triton X-100, 1 mM sodium orthovanadate, 1μM okadaic acid, 5 mM sodium pyrophosphate, 30 mM β-glycerophosphate, 5 mM EDTA, protease inhibitors (Complete, Roche, Cat. No 11697498001)). Homogenates were centrifuged at 15000 rpm for 15 min at 4°C. The resulting supernatant was collected, and protein content determined by Quick Start Bradford Protein Assay (Bio-Rad, 500-0203). 20 μg of total protein were electrophoresed on 10% SDS-polyacrylamide gel, transferred to a nitrocellulose blotting membrane (Amersham Protran 0.45 μm, GE Healthcare Life Sciences, 10600002) and blocked in TBS-T (150 mM NaCl, 20 mM Tris–HCl, pH 7.5, 0.1% Tween 20) supplemented with 5% non-fat dry milk. Membranes were incubated overnight at 4°C with the primary antibody in TBS-T supplemented with 5% non-fat dry milk, washed with TBS-T and next incubated with HRP-conjugated anti-mouse IgG (1:2000, DAKO, P0447) or anti-rabbit IgG (1:2000, DAKO, P0448) and developed using the ECL detection kit (PerkinElmer, NEL105001EA).

#### Antibodies

Rabbit anti-CCDC88C (1:500, Sigma, HPA005832), rabbit anti-KCTD17 (1:500, Thermo Fisher, PA5-72101), mouse anti-SYNJ1 (1:500, Thermo Fisher, MA3-936), rabbit anti-TRPM7 (1:1000, Novus Biologicals, NBP2-20739), rabbit anti-SLC9A5 (1:500, Abcam, ab191528), rabbit anti-TIA1 (1:1000, Abcam, ab40693), rabbit anti-U2AF2 (1:500, Santa Cruz, sc-53942), mouse anti-RBFOX1 (1:2000 for mouse and 1:1000 for human samples, Merk Millipore, MABE985), mouse anti-RBFOX2 (1:500, Abcam, ab57154), rabbit anti-RBFOX3 (1:1000, Merk Millipore, MAB377), rabbit anti-ELAVL4 (1:500, Abcam, ab96474), rabbit anti-ELAVL2 (1:500, Abcam, ab96471), rabbit anti-ELAVL1 (1:500, Abcam, ab200342), rabbit anti-HNRNPC (1:500, Abcam, ab10294), rabbit anti-ELAVL3 (1:500, Abcam, ab129254), rabbit anti-PTBP1 (1:1000, Abcam, ab5642), rabbit anti-PTBP2 (1:2000, Merk Millipore, ABE431), mouse anti-β-ACTIN (1:25000, Sigma, A2228), rabbit anti-VINCULIN(1:20000, DAKO, P0448).

#### Motif enrichment analysis

A total of 73 orthologous exons with differential splicing in both human and mouse were evaluated in parallel for RBPs motif enrichment. As a control set, we selected 914 conserved AS exons with no detectable changes (|ΔPSI|<5) in the RNA-seq experiments of both species. RBP motifs deposited in the CIS-BP database (http://cisbp-rna.ccbr.utoronto.ca/index.php; downloaded july 2016) were scanned separately in exons, 250 intronic bases upstream the exon (3’ splicing signal – 3’ss, excluding 20bp within the 3’ splice site) and 250 intronic bases downstream the exon (5’splicing signal – 5’ss, excluding 6bp within the 5’ splice site). Motif occurrences (log-odd ≥6 for the respective RBP Position Weight Matrix) for each RBP were counted for regulated and control exons respectively. A motif can be counted zero or once in a given sequence. Motif enrichment for each RBP in regulated AS exons compared to the control set was evaluated using Fisher’s exact test. To correct for multiple testing, Benjamini-Hochberg FDR was used for adjustment of p-values. Enrichment was considered significant when the adjusted P < 0.01.

#### Data analysis

Statistical analysis was performed with SPSS 21.0 (SPSS® Statistic IBM®). Data are represented as Mean ± SEM (Standard Error of the Mean). The normality of the data was analyzed by Shapiro-Wilk. For two-group comparison, two-tailed Student’s t-test was performed (data with normal distribution) or Mann-Whitney U-test (data with non-normal distribution) was performed. A critical value for significance of *P* < 0.05 was used throughout the study. Benjamini-Hochberg correction was applied for multiple testing in RNA-seq analysis and motif enrichment analysis, were enrichment was considered significant when the adjusted P < 0.01.

## SUPPLEMENTARY FIGURE AND TABLE LEGENDS

**Supplementary Fig. 1.**
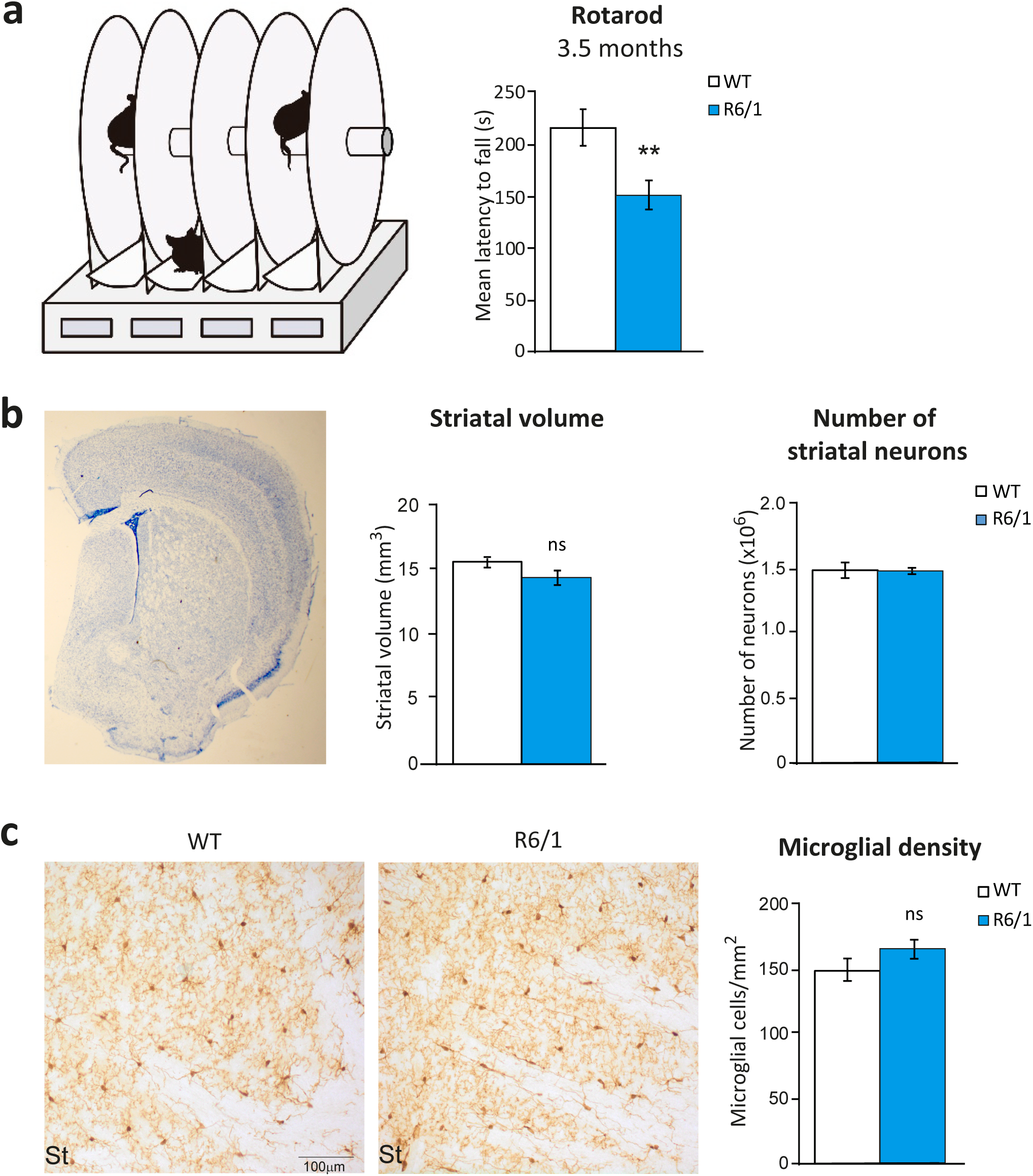
**(a)** Rotarod test. Mean latency to fall from the rod in the four accelerating testing trials of WT (*n*=9) and R6/1 (*n*=11) mice at 3.5 months of age. (Student’s t-test; *\*\* P < 0*.*01*). Data represent mean ± SEM. **(b)** Representative image of toluidine blue staining in coronal sections from R6/1 mice and quantification of the number of neurons and the striatal volume measured in coronal sections of WT (*n*=3) and R6/1 (*n*=3) mice at 3.5 months of age. (Student’s t-test). Data represent mean ± SEM. **(c)** Representative images of Iba1 immunostaining in sagittal sections from R6/1 (*n*=3) and WT (*n*=3) mice at 3.5 months of age and quantification of the density of microglial cells. (Student’s t-test). Data represent mean ± SEM.

**Supplementary Fig. 2.**
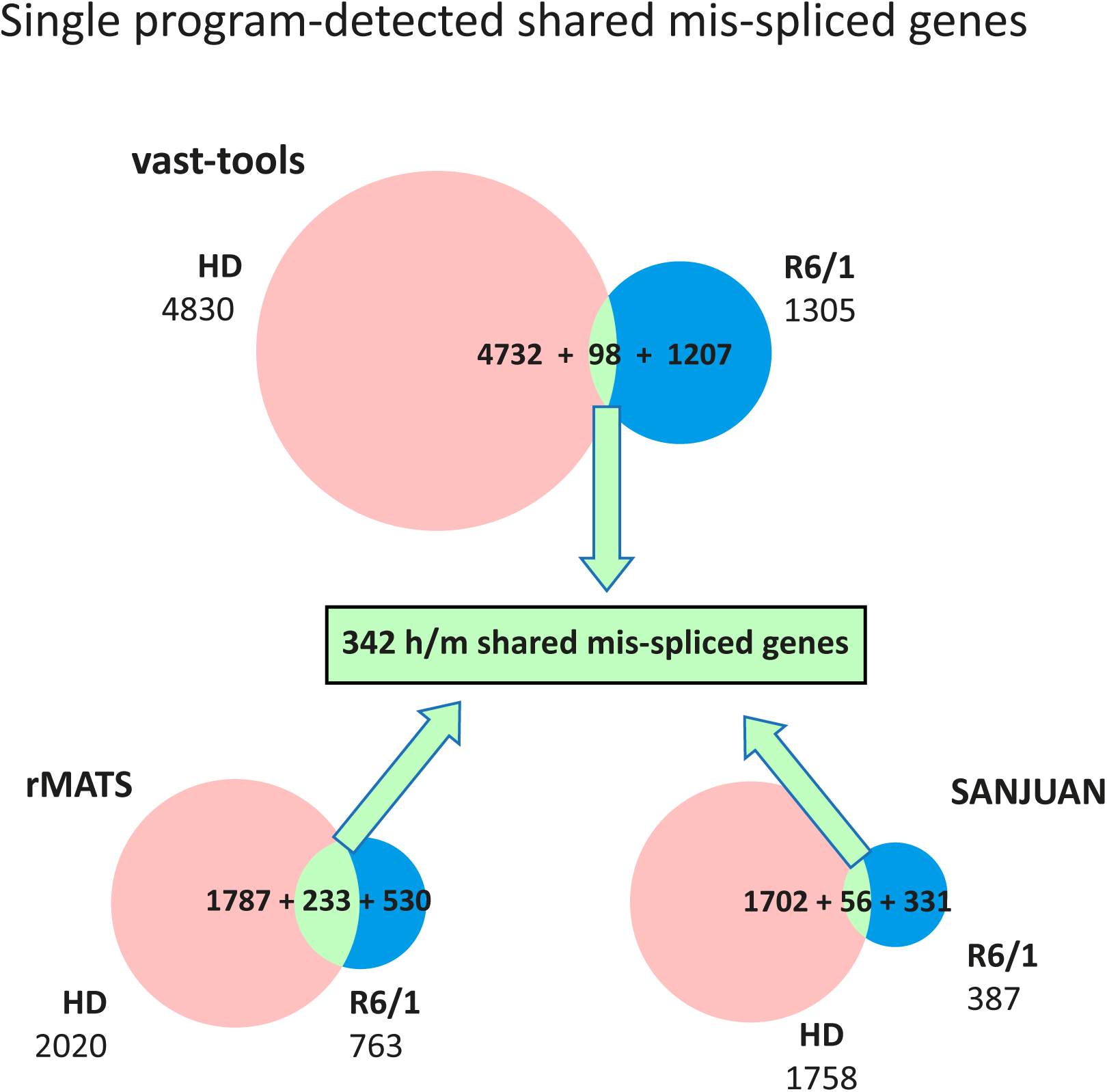
Venn diagrams showing the number of mis-spliced genes in striatum of HD patients and R6/1 mice with respect to controls and those shared by both species, according to each of the AS-analysis tools.

**Supplementary Fig. 3.**
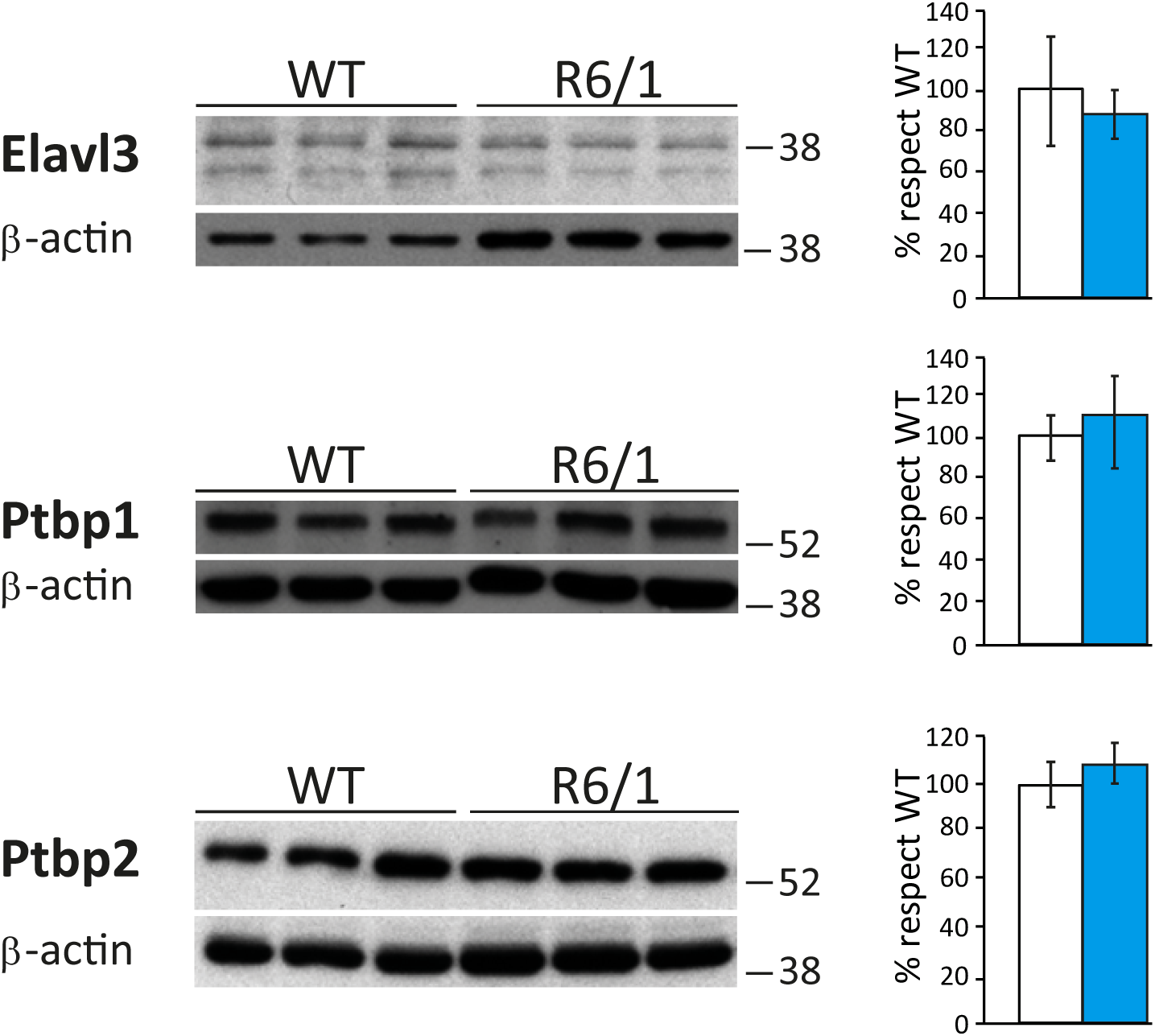
Elavl3, Ptbp1 and Ptbp2 levels in WT and R6/1 striatum and quantification normalized with β-actin (*n*=7-12). (Student’s t-test). Data represent mean ± SEM.

